# Moving to the city changed you! Rapid photobiont turnover enables acclimation of lichen symbioses to urban environments

**DOI:** 10.64898/2026.08.14.744966

**Authors:** Natália Mossmann Koch, Laima Liulevičius, Abigail Meyer, Anja Nilles, Lindsey Kemmerling, Emilie Snell-Rood, Daniel Stanton

## Abstract

Symbioses are widespread and highly successful but vulnerable to the stress sensitivity of either symbiont. In some symbioses, turnover of symbionts has been shown to confer resilience to stressors. While similar mechanisms have been proposed for lichen symbioses, direct evidence for rapid adaptive symbiont turnover has not been shown. We tested the photobiont community composition and physiological responses of the foliose lichen symbiosis *Flavoparmelia caperata-Trebouxia* to urbanization-induced stress in a transplant experiment. We found evidence for significant compositional change in the photobiont community along an urbanization gradient (measured as vegetation cover), reflecting a turnover in dominance of *Trebouxia* OTUs from A46 to I05 in more urbanized transplant sites. This change in symbiont composition is associated with a greater physiological tolerance for urbanization, consistent with the hypothesized adaptive role of photobiont turnover. These findings support rapid photobiont turnover as a potential adaptive response to environmental change in lichen symbioses.

## Introduction

Decades of research highlights the importance of mutualistic symbioses across ecosystems (Kiers *et al*. 2010; Rolshausen *et al*. 2020). However, the multispecies nature of mutualisms creates opportunities for both rapid adaptation and niche expansion (Kiers *et al*. 2010; Nathan *et al*. 2023) and potential vulnerabilities to environmental change (Cruz *et al*. 2023; Kiers *et al*. 2010). At the association scale, the turnover of individual genotypes within a symbiosis can provide a mechanism for rapid acclimation in response to sudden or gradual environmental changes. Notably, in the coral-dinoflagellate symbiosis, recent work has shown that *Symbiodinium* communities change following marine heat waves (Van Nynatten *et al*. 2025) and over time and space (Grillo *et al*. 2025). This persistence of the symbiotic whole through changes in the parts (“it’s the song not the singers” *sensu* Doolittle & Booth (2017)) may be a key source of symbiotic resilience and flexibility, however, nonetheless requires both availability of suitable symbionts and the capacity for turnover on the timescales of the triggering change. While this adaptive ability has been shown, within constraints, for corals (i.e. changes in population densities and pigment concentrations, Buddemeier et al. 2004; Cavailles et al. 2025), and Rhizobia (i.e. preference for the “best” strain available, Westhoek *et al*. 2021), it remains less certain for other iconic symbioses such as lichen-forming associations of fungi and photoautotrophs.

In lichen symbioses, the turnover of symbionts, particularly of algal partners (photobionts), has been suggested as a mechanism for environmental adaptation (Evankow *et al*. 2025; Spribille *et al*. 2022; Stanton *et al*. 2023; Williams *et al*. 2017). This potential is strengthened by the occurrence of not only algal genotypic diversity (Dědková *et al*. 2025; Gasulla *et al*. 2025; Piercey-Normore 2006) but also physiological/functional diversity (Casano *et al*. 2011; Schofield *et al*. 2003) within individual lichen thalli. Within the lifespan of a lichen, photobiont community diversity can increase, suggesting the potential for continued incorporation of new photobiont genotypes (Kono *et al*. 2025). However, although there is clear evidence for turnover of dominant photobionts over spatial environmental gradients, such as climate, elevation and nitrogen pollution (Dal Grande *et al*. 2017; Gasulla *et al*. 2025; Medeiros *et al*. 2021; Rolshausen *et al*. 2020; Werth & Sork 2014), evidence for temporal turnover on ecologically relevant timescales remains less clear. A long-distance transplant experiment with the terricolous *Psora decipiens*-*Myrmecia* association did not find evidence for incorporation of new algae (Williams *et al*. 2017) while a short-distance transplant of the epiphytic *Evernia mesomorpha*-*Trebouxia* association across a temperature gradient found directional changes in photobiont communities, but in a context of extensive thallus mortality (Meyer *et al*. 2023).

Urban environments are affected by both regional and local scale changes on climate, most notably the urban heat island (UHI) effect (Landsberg 2011; Li *et al*. 2024), combined with a suite of other stressors, including air pollution. Due to this, they can be used as proxies to test the effects of environmental changes on sensitive symbiotic associations, such as lichens. Lichen communities have been used as indicators of the urban environment for more than a hundred years (Nylander 1866; Sernander 1926). While the majority of studies of lichen ecology in urban contexts have focused on community scale responses to pollutants (Koch *et al*. 2019; Rocha *et al*. 2022; Schram *et al*. 2015), or the accumulation of pollutants in thalli as indicators of air quality (Dupont *et al*. 2025; Koch *et al*. 2018; McCarthy *et al*. 2009; Van der Wat & Forbes 2015), surviving urban lichen thalli can also show physiological stress responses to urbanization (Sujetovienė & Galinytė 2016). Pollutants such as lead, cadmium, chromium and NO_X_ can reduce chlorophyll content (e.g. Karakoti *et al*. 2014; Piccotto *et al*. 2011) or photosynthetic capacity (e.g. Styburski & Skubała 2023). However, with improving air quality in recent decades in many urban areas, there is growing evidence for urban microclimates becoming the primary driver of urban lichen communities (Counoy et al. 2025; Koch et al. 2019; Munzi et al. 2014).

Increased impervious surface area and decreased area of greenspaces are features of the urban landscape that typically result in increased temperatures and lower humidity (Pickett *et al*. 2011). With less vegetation, there is consequently an increase in the urban Vapor Pressure Deficit (VPD) causing faster loss of humidity by lichen communities, which rely greatly on vapor hydration. All these factors make NDVI (Normalized Difference Vegetation Index) a useful proxy of urbanization (Ju *et al*. 2024; Martinez & Labib 2023).

Most studies showing an effect of urbanization on lichen symbioses are focused on community changes (Claerhout *et al*. 2026; Käffer *et al*. 2011; Varela *et al*. 2018), or photobiont physiology (Piccotto *et al*. 2011; Tarhanen *et al*. 2000). Microbial communities can also shift rapidly in lichen thalli transplanted to higher temperatures (Yang *et al*. 2026). However, photobiont identity may play an underappreciated role in determining lichen sensitivity to urban environments. More recent work showed that common lichen-forming algal genera such as *Trebouxia* can exhibit considerable variability in their tolerance to urban stressors among strains (Gasulla *et al*. 2025), but a clear connection between temporal environmental changes and turnover of the lichen photobiont community has not been shown.

To experimentally evaluate the effects of urban conditions on lichen photobiont identity and physiology, we conducted a series of rural to urban transplants across an urbanization gradient using the widespread temperate *Flavoparmelia caperata-Trebouxia* lichen association. We used vegetation cover to define our urban gradient, measured as NDVI. We evaluated impacts on photobiont community composition as well as the corresponding physiological performance of the lichen thalli. We predict that in lichen thalli transplanted into more urbanized sites, photobiont community composition will shift from showing dominance of clades that are less stress tolerant to clades that are more stress tolerant. This shifted composition will be reflected in the physiological performance (carbon assimilation and respiration, chlorophyll fluorescence) at the thallus level.

## Material and Methods

### Sampling the monitoring species and experimental design

Thalli of the lichen association *Flavoparmelia caperata-Trebouxia,* commonly known as the “Common Green Shield Lichen”, were collected with their bark substrate from fallen trees at Cedar Creek Ecosystem Science Reserve (45°24’ 04.0”N, 93° 11 ‘56.0”W). All samples were taken from the same source population, as represented by a single stand of deciduous forest. From this initial population, two samples were used as the baseline in the first year (2022), and eight samples were collected and tested in the second year (2023).

*Flavoparmelia caperata* (L.) Hale is a classic species of Parmeliaceae (Ascomycota) used in biomonitoring studies (Expósito *et al*. 2020; Will-Wolf *et al*. 2017). It is a widespread foliose lichen associated with unicellular algae from the genus *Trebouxia* (Trebouxiophyceae, Chlorophyta) as its photobiont–the best studied lichenized algae so far. This lichen species is also somewhat tolerant to air pollution, including metals, sulfur and nitrogen-rich deposition (Chahloul *et al*. 2023, 2024; Will-Wolf *et al*. 2017).

After 48h air-dry acclimation to laboratory conditions (23°C, ∼40%RH) all thalli were exposed to high humidity until hydrated, tested for baseline chlorophyll fluorescence and photographed (see details below). Thalli were then tied to plastic mesh (30 x 20cm), and the transplants were placed across the metropolitan area of the Twin Cities (Minneapolis, Saint Paul and surrounding cities), Minnesota, USA. At each of 29 unique sites, one or two transplant replicates were deployed, and replicates were deployed at the same time at Cedar Creek (the original population source site) as a control. Part of these transplants were deployed for a total of 10 months in 2022 (19 sites) and 6 months in 2023 (10 sites and the three control replicates). Sites were selected to represent a wide gradient in urbanization and air pollution exposure, including coinciding with sites of another ecological research in the Minneapolis-Saint Paul LTER site (see Figure 1). For ecophysiological, elemental and photobiont composition analyses, transplanted samples were collected at 5 and 10 months (2022) and at 6 months (2023).

**Figure 1.**
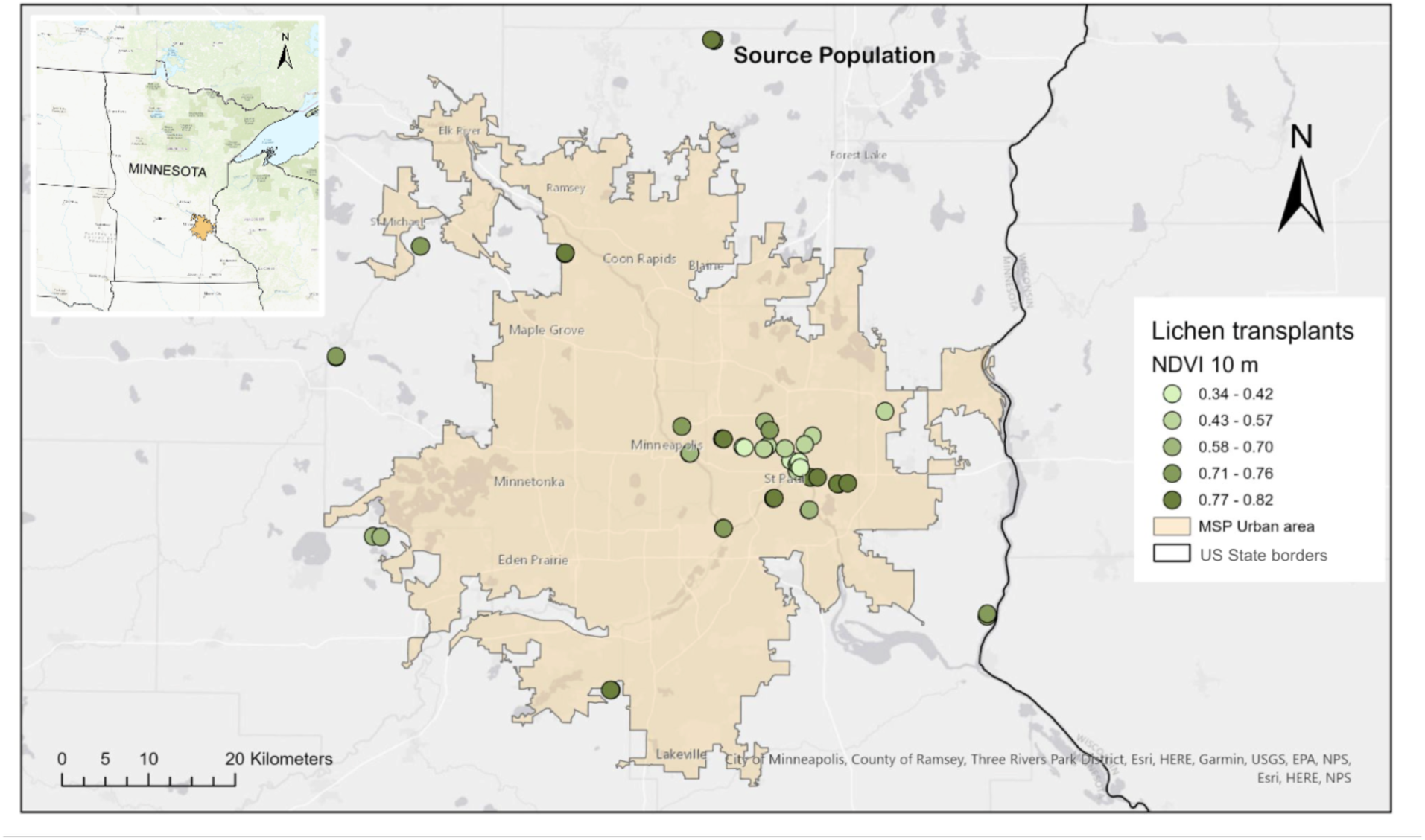
Map of the study location. The urban area as defined by the US Census Bureau of the Twin Cities (Minneapolis and St. Paul) in Minnesota, USA is colored in orange. Circles in shades of green mark lichen transplant locations shaded by NDVI value at a resolution of 10 m. Darker circles indicate higher values of NDVI. North of the urban area is the source population, sampled at Cedar Creek Ecosystem Science Reserve. The inset shows the study area within the state of Minnesota.

### Characterizing the urbanization gradient

We used NDVI as a proxy for urbanization (Du *et al*. 2019; Weng *et al*. 2004). This metric showed positive correlation with higher cover of trees, shrubs and grassland in the urban environment (Martinez & Labib 2023), while being negatively correlated with LST (Land Surface Temperature) in warmer months (Sun & Kafatos 2007; Weng *et al*. 2004) and used for remote sensing of the Urban Heat Island Effect (UHI) (Zhang *et al*. 2012). At the same time, LST is known to be positively correlated with impervious surface cover (Yuan & Bauer 2007).

We calculated the average NDVI for each tree on which lichens were transplanted at the radius of 10 m, the smallest resolution available using the terra package in RStudio (Hijmans *et al*. 2026). The NDVI layer used was acquired in 2018 and derived from Landsat satellite imagery processed via Google Earth Engine by the Natural Resources Research Institute, University of Minnesota Duluth (Johnson *et al*. 2022). We also extracted data on LST at the same resolution (10 m). Data on LST was published by the Metropolitan Council of the Twin Cities Area (Metropolitan Council 2022) and derives from measurements taken on September 1, 2022, using satellite imagery from Landsat 9 and downscaled to 10 meters using Copernicus Sentinel-2. This fine spatial scale was selected since lichens are sensitive to microclimatic conditions (Stanton *et al*. 2023). For our sites, NDVI and LST showed strong correlation (⍴ = - 0.84, P < 0.0001), so only NDVI was used for subsequent analyses. We created a map of our study sites overlaid with NDVI in ArcGIS Pro version 3.3 (Esri Inc. 2024), along with the boundary of the Twin Cities metro area that is designated as an Urban Area (US Census Bureau) (Figure 1).

### Ecophysiological analyses

Ecophysiological traits related to carbon balance and photosynthesis, namely chlorophyll fluorescence, carbon assimilation and respiration rates, were measured on lichen thalli to test their response to transplant site environments.

Prior to measurement, thalli taken from the field were placed on the lab bench (23°C, ∼40% RH) until they air dried and equilibrated with room temperature (at least 24 hours). A subsample (around 2 cm^2^) was then removed from a healthy, growing lobe and carefully detached from underlying bark without damage. To achieve vapor hydration, thalli were placed in sealed containers (∼700 cm^3^) over distilled water for 18 hours at low light (<15 μmol m^-2^ s^-1^). These conditions achieved complete equilibration with humid air within hours (Meyer *et al*. 2024; Phinney *et al*. 2018) and mimicked natural nocturnal conditions.

After vapor hydration, thalli were liquid hydrated with a fine spray of distilled water applied progressively to enable consistent, gentle hydration (sprayed directly three times every 2 minutes, for 20 minutes, turning the specimen to hydrate both sides equally). Thalli were allowed to drain excess water between sprays, to obtain complete hydration without over-saturation (Meyer *et al*. 2024).

#### Chlorophyll fluorescence - Fv/Fm

To test maximum photosynthetic efficiency of PSII (Fv/Fm), each specimen (n = 84) was tested under liquid activated conditions and chlorophyll fluorescence was measured using a red-light IMAGING-PAM m-series chlorophyll fluorometer (Walz, Effeltrich, Germany). Each thallus was placed in a 100% humidity chamber, and a saturating pulse was applied after 20 minutes of dark acclimation to record Fv/Fm, which was automatically calculated in ImagingWinGigE version 2.56p.

#### Assimilation and respiration rates - CO_2_ exchange

Thallus assimilation (CO_2_ acquisition) and respiration (CO_2_ release) rates were measured on the majority of the same thalli (n = 68) tested for chlorophyll fluorescence using a portable Infra-Red Gas Analyzer (LI-6800, LI-COR, Lincoln, NE, USA) with the Aquatic Chamber attached to it (6800-18, internal volume of 20 cm^3^). To reduce rates of water loss during measurement (and thus maintain constant hydration levels over the course of measurement periods) high humidity conditions were applied: RH target 95% (realized RH ∼91%), air temperature ∼26 °C in the chamber, 500 µmol/s flow rate and reference CO_2_ of 410 ppm. For light measurements, the head light was set at a PAR of 1000 μmol m^-2^ s^-1^ and the color ratio of r90b10. Gas-exchange measurements were recorded following stabilization of IRGA readings for at least two minutes, at which point 10 replicate measurements at 1-2 second intervals were logged to obtain a mean value. Dark measurements always preceded light measurements. Gross assimilation, representing the carbon uptake potential of the photobiont, was calculated by subtracting the dark measurements (respiration) from the light measurements (net assimilation).

### Heavy metal analyses

Air pollution can also impact urban lichen physiology (Backor & Loppi 2009; Sujetovienė & Galinytė 2016). To account for air quality as a potential conflating factor, we also measured the elemental content of thalli to assess the accumulation of aerial deposition over time. A subsample of ∼10 mg of lichen tissue was sampled from each transplant (n = 88) within no more than two weeks after their field retrieval and sent to QBIC (Northwestern University - Quantitative Bulk-Elemental Information Core) for ICP-MS analysis. Samples were digested in acid prior to running and run using a quadrupole-based inductively coupled plasma mass spectrometer for the detection of most elements in the low ppb concentration range (usually 0.02-10,000 ppb). Data was obtained for 18 elements (Na, Mg, Al, P, K, Ca, V, Cr, Mn, Fe, Co, Ni, Cu, Zn, As, Se, Cd, Pb), however, our further analyses were focused on the elements with known detrimental effects for the lichen photobiont (Fe, Pb and Zn), based on the post-transplant metal accumulation, and the previous accumulation was measured from the source population.

### Photobiont extraction and sequencing

Photobiont composition in the transplanted thalli was assessed using amplicon sequencing targeting the photobiont genus *Trebouxia* following the methods of Meyer et al. (2023). Briefly, 0.03 g of lichen tissue was taken from a healthy growing edge of each thallus, (2022: source n= 2, 5m transplant n= 30, and 10m transplant n= 25; 2023: source n= 8, 6m transplant n= 22). Because *Trebouxia* DNA is considered to be stable (Honegger 2003), DNA extractions were done with fresh thalli (or kept at -20℃ for no longer than 3 months) and performed following a modified CTAB (cetyltrimethylammonium bromide) extraction protocol. Following DNA extraction, a two-step PCR amplification protocol was used with the primers ITS1T (GGA AGG ATC ATT GAA TCT ATC GT) and ITS2T (TTC GCT GCG TTC TTC ATC GTT) (Kroken & Taylor 2000) which were modified for Illumina sequencing (Smith & Peay 2014).

PCR reactions were performed using Phusion HF buffer and Phusion taq polymerase (New England Biolabs, Ipswich, Massachusetts, USA) and the reactions were carried out using a touchdown-PCR protocol in a Veriti 96-Well Thermal Cycler (Applied Biosystems, Foster City, California, USA). Final PCR products were cleaned using either the Mag-Bind TotalPure NGS Kit (Omega Bio-tek, Norcross, Georgia, USA) or Just-a-Plate 96 PCR Purification and Normalization Kit (Charm Biotech, Cape Girardeau, Missouri, USA) and quantified using the Qubit hs-DS-DNA kit on a Qubit 2.0 Fluorometer (Invitrogen, Carlsbad, California, USA). 4 nM of each sample was then pooled and submitted to the University of Minnesota Genomics Center (UMGC) for sequencing on an Aviti Freestyle Medium sequencer (Element Biosciences, San Diego, California, USA).

### Sequence Analytics

Raw, demultiplexed amplicon data were obtained from the University of Minnesota Genomics Center and processed using QIIME2 version 2024.10 (Bolyen *et al*. 2019). Primer sequences were trimmed from R1 reads using the Cutadapt plugin (Martin 2011). To trim low quality reads (PHRED score <20), the DADA2 plugin (Callahan *et al*. 2016) was used to truncate sequences to 274 base pairs. The VSEARCH plugin (Rognes *et al*. 2016) was used for open-reference clustering, grouping sequences into OTUs based on 97% similarity to a comprehensive *Trebouxia* reference dataset (Muggia *et al*. 2020). Following (Reitmeier *et al*. 2021), OTUs that represented less than 0.10% relative abundance (441) were removed. An OTU table was exported for downstream analysis in R version 4.3.1 (R Core Team 2020).

### Statistical Analyses

We assessed the overall trend of OTU richness per lichen thallus along the urbanization gradient with a linear model (simple linear regression using “lm” function in R) of richness as a function of increasing NDVI.

Because two algal genotypes (A46 and I05) were much more widespread and dominant than others (>70% of reads in 65/88 thalli - Fig. 2), many of the further analyses focused on those two genotypes. To model photobiont composition along the NDVI sampling gradient, two modeling approaches were used. The first modeled continuous proportions of the two most dominant genotypes, A46 and I05, using a beta-binomial regression that accounts for overdispersion. For this model, A46 and I05 reads were rarefied and a generalized linear model was constructed with the glmmTMB package (McGillycuddy *et al*. 2025) using family = betabinomial. The second approach modeled photobiont dominance as a binary outcome whereby any sample where the proportion of A46 or I05 reads was >70% was labeled as dominated by that OTU, respectively. Six samples were lost due to lack of I05 or A46 dominance (proportion < 70%). A logistic regression was fitted where we modeled the probability of A46 dominance as a function of NDVI with the stats package (R Core Team 2020) using family = binomial. Model diagnostics were evaluated with the DHARma package (Hartig 2016). The estimated dispersion parameter (theta = 0.42) indicated that the data is overdispersed, warranting the beta-binomial modelling approach.

**Figure 2.**
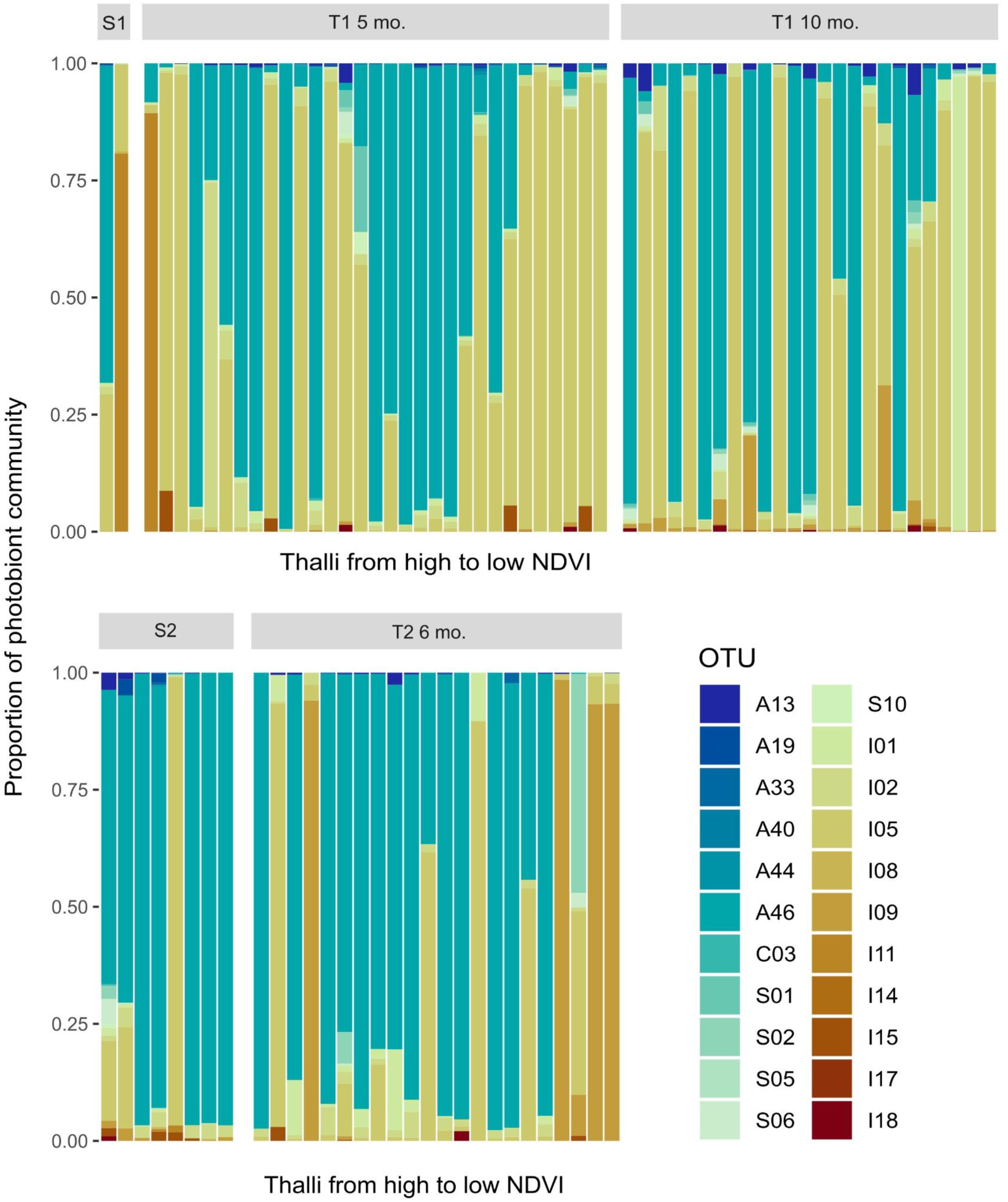
Photobiont composition of *Trebouxia* OTUs (20 total) in each *Flavoparmelia caperata* thalli along an urbanization gradient, arranged from high to low NDVI (10 m resolution). Transplant experiment 1 was conducted in 2022: S1 is source population 1, T1 are transplants, which were retrieved after 5 months and 10 months (T1 5 mo. and T1 10m). Transplant experiment 2 was conducted in 2023: S2 is source population 2, and T2 are transplants which were retrieved after 6 months (T2 6 mo.).

We also analyzed changes in the ecophysiological performance of lichens transplanted along the NDVI gradient, including the source site, as well as individuals of the source population at time of first collection. To better understand the relationship of lichen photobiont algal clade composition and thallus ecophysiology, we used the same threshold of 70% or more reads belonging to one OTU and then assigned that OTU as the site dominant. We modeled each parameter (gross assimilation, net assimilation, respiration, and Fv/Fm) as a response to increasing NDVI interacting with OTU dominance in a linear model (least-squares linear regression, ‘lm’ function in R) fitted separately to each OTU, as well as a response to changes in heavy metals content. We also tested the overall correlation of each ecophysiological parameter with increasing NDVI without separating thalli into photobiont OTUs using the function ‘cor.test’ with Spearman as the correlation method.

Statistical analyses were performed in RStudio (version 4.5.2) (RStudio Team 2025), and data was visualized using ggplot2 (Wickham 2016).

## Results

### Clades turnover led by the urban microclimate

Urbanization led to shifts in photobiont dominance with increasing NDVI within the lichen thalli, A46 and I05 being the dominant OTUs of *Trebouxia* (Figures 2 and 3, Table S1) present in all the 88 tested specimens (above 70%). A total of 22 OTUs were found in the sampled thalli and the richness of algal OTUs did not significantly linearly increase or decrease as a response to decreasing NDVI (R^2^adj = 0.01, P = 0.15). Median NDVI for I05 dominated thalli was 0.57 (IQR: 0.43, 0.77) and for A46 thalli was 0.76 (IQR: 0.62, 0.79).

**Figure 3.**
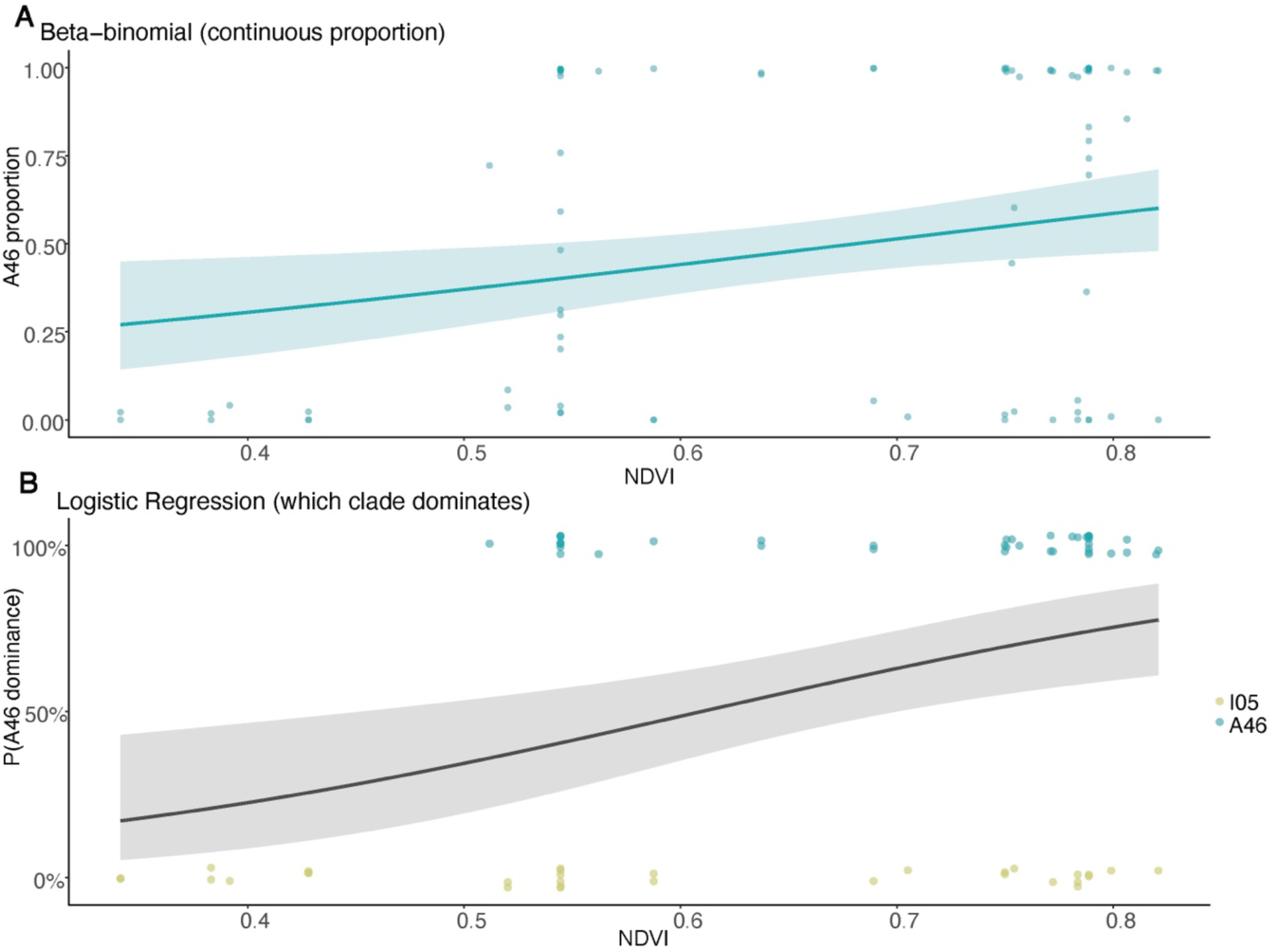
Shifts of the photobiont composition in *Flavoparmelia caperata* with increasing NDVI (A) Beta-binomial regression of the proportion of A46:I05 against NDVI, capturing how photobiont community proportions shift with increasing NDVI. (B) Logistic regression of the probability of A46 dominance against NDVI, capturing how thalli dominated by A46 versus I05 shift with increasing NDVI. Lines represent fitted values for each model and 95% CI.

The beta-binomial regression showed a significant positive relationship between increasing NDVI and proportion of A46 reads in a lichen transplant (B=2.98 +/- 1.17 SE, z = 2.54, p = 0.01), or more simply, for every 0.1 unit increase in NDVI, the odds that a given sequencing read was A46 increased by a factor of 1.35 (Figure 3a). The logistic regression model showed the same pattern: higher NDVI increased the probability of a lichen thallus being dominated by A46 (B=6.00 +/- 1.92 SE, z = 3.12, p = 0.002; Figure 3b). The logistic model predicted that for every 0.1 unit increase in NDVI, the odds of a lichen thallus being A46 dominant increased by a factor of 1.8.

There was a significant effect of urbanization on the ecophysiological parameters tested on *Flavoparmelia caperata* thalli, with responses varying by dominant photobiont (Figure 4). Overall, gross and net assimilation (ρ = 0.28; P = 0.04; and ρ = 0.27; P = 0.05, respectively), as well as Fv/Fm (ρ = 0.45; P = 0.0001) were positively correlated with NDVI, but not respiration (ρ = 0.03; P = 0.82). Thalli dominated (frequency > 70%) by photobiont I05 did not show a physiological response to transplant site NDVI (Gross assimilation - Estimate = 1.21, SE = 3.29, t-test = 0.37, P = 0.72; Net assimilation - Estimate = 0.66, SE = 3.68, t-test = 0.18, P = 0.86; Respiration - Estimate = -0.55, SE = 0.81, t-test = -0.68, P = 0.51; Fv/Fm - Estimate = 0.10, SE = 0.14, t-test = 0.67, P = 0.51; Table S1 and Figure 4), whereas A46 exhibited reduced Fv/Fm in low NDVI sites (Estimate = 0.45, SE = 0.12, t-test = 3.75, P = 0.0007) and the same pattern was observed for gross (Estimate = 7.30, SE = 3.09, t-test = 2.37, P = 0.03) and net assimilation (Estimate = 8.51, SE = 3.29, t-test = 2.59, P = 0.02), but not for respiration (Estimate = 1.20, SE = 1.55, t-test = 0.78, P = 0.45) as showed in Figure 4 and Table S1.

**Figure 4.**
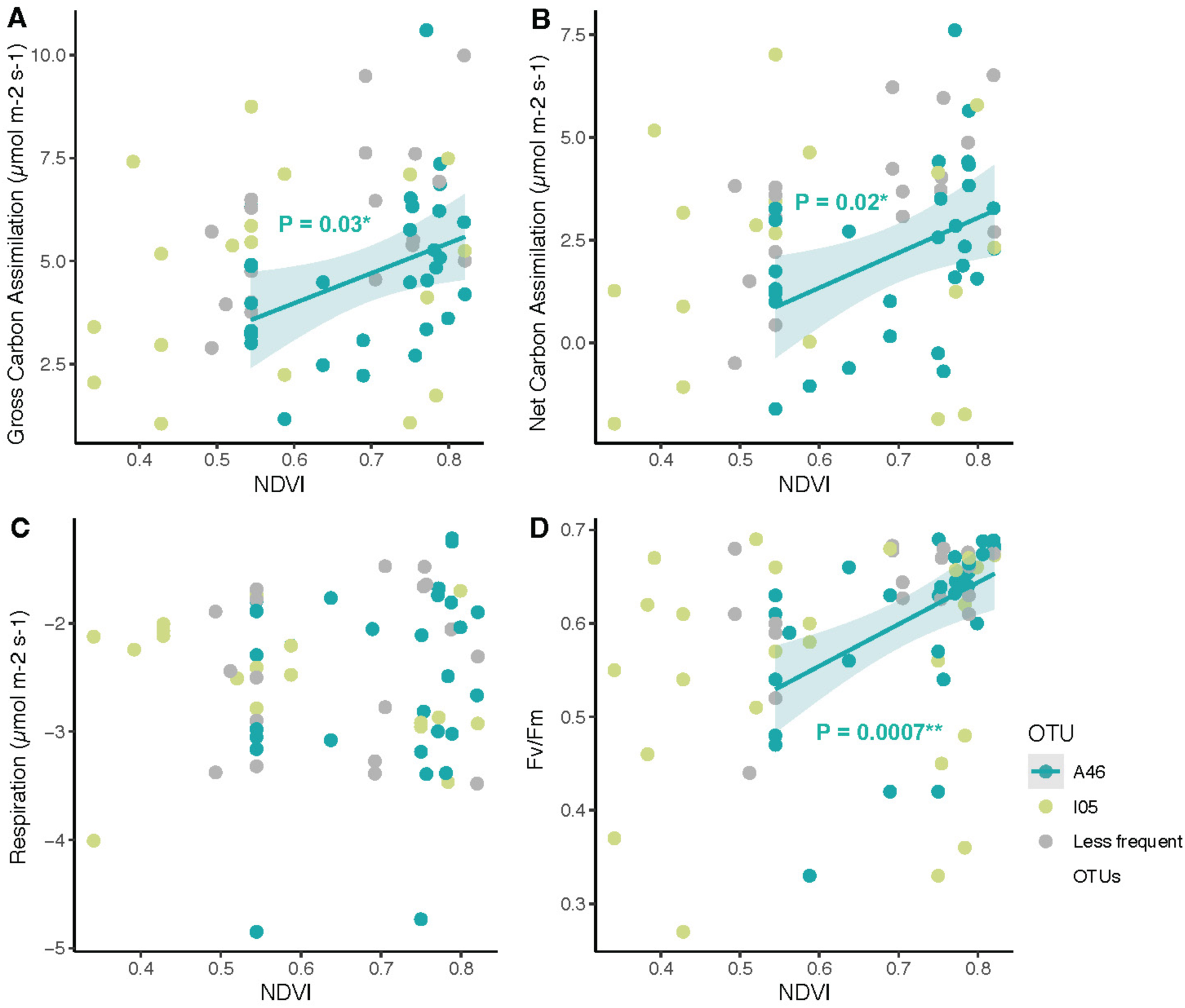
Physiological performance metrics of *Flavoparmelia caperata* thalli according to the OTU/Clade that predominates in their photobiont community (>70% A46 or I05) with increasing NDVI. Points depict individual thalli that are colored based on this dominance. Metrics include **A.** gross carbon assimilation (µmol m^-2^ s^-1^), **B.** net carbon assimilation (µmol m^-2^ s^-1^), **C.** respiration (µmol m^-2^ s^-1^), and **D.** maximum photosynthetic efficiency of PSII (Fv/Fm).

### Heavy metals

We only found significant but weak relationships between the Zinc content and respiration and photosynthetic potential (Fv/Fm), where increasing Zn was related to lower Fv/Fm in the transplanted thalli dominated by I05 (Estimate = -0.004, SE = 0.002, t-test = -2.22, P = 0.04). We did not find significant patterns for the deposition of other heavy metal and lichen performance (Supplement Figure S1).

## Discussion

We found evidence for significant and rapid directional shifts in the composition of algal communities of the *Flavoparmelia caperata* lichen symbiosis in response to exposure to urbanization. These changes primarily take the form of a change in dominant genotypes of *Trebouxia* from A46 to I05 within less than a year, when comparing the transplanted specimens with their source population. We interpret shifts in community composition as turnover in the photobiont population because they appear in the growing edges, across multiple independent thalli and transplant sites. Because thalli dominated by A46 showed some physiological sensitivity to urbanization (significant decrease in assimilation and photosynthetic efficiency), this genotypic turnover may contribute to buffering the environmental effects on carbon uptake, providing a mechanism for thallus-level homeostasis across environments (Stanton et al. 2023).

Shifts in lichen algal communities have been documented across climatic gradients for numerous taxa at this point, from geographically short but environmentally steep mountain gradients (Dal Grande *et al*. 2017, 2018) to large regional and global patterns (Rolshausen *et al*. 2020). These patterns, however, mostly reflect long-established environmental gradients. By using large-distributed transplant experiment, we are now able to show that photobiont communities can shift composition on relatively rapid timescales of months to years, which is especially striking considering the slow growth rates associated with lichens, including in urban settings (1-5 mm yr-1, Armstrong & Bradwell 2011; Kubo & Ohmura 2025). The algal layer of lichen thalli has previously been shown to be dynamic in thickness on seasonal (Tretiach *et al*. 2013) to multi-annual (e.g. Johansson *et al*. 2011) scales, however this may be the first evidence of rapid genotypic change of photobionts consistent with within-thallus environmental filtering.

The observed changes in algal community composition may have arisen through multiple mechanisms: acquisition of new algal partners from the environment, changes in existing algal partners through selective loss of algal cells and/or differences in algal genotype growth.

While acquisition of novel algal symbionts has been well documented for some lichen associations on longer time scales (Dědková *et al*. 2025), the community composition changes observed in our transplants largely reflect genotypes present in the source population (Fig. 2). Therefore, while acquisition of I05 algal cells is possible, changes in the existing population would be the more parsimonious interpretation. The changes in algal communities are also unlikely to be due solely to selective death of the A46 photobiont: many of the transplanted thalli appeared physiologically healthy, including showing levels of chlorophyll fluorescence and carbon fixation comparable to starting conditions and source populations. This relative physiological stasis is not consistent with solely the loss of a previously dominant algal genotype, but rather also requires the compensatory growth of I05 populations.

This contrasts with a previously documented algal community change in *Evernia mesomorpha*, in which the poor health of the higher temperature transplants made it difficult to distinguish between selective survival and selective growth of the photobionts (Meyer *et al*. 2023). These considerations would suggest that the change in composition most likely arises from differential growth within the thallus, favoring I05 genotypes relative to other genotypes in more urbanized settings. *Trebouxia* strains have been shown to differ in their physiology, including across urbanization gradients (Gasulla *et al*. 2025). We found similar differences between *Trebouxia* strains, with a significant negative correlation between carbon uptake and urbanization in thalli dominated by one common strain (A46) but not the other (I05) (Fig 4). While further studies of strain-specific physiological tolerances (especially *in talo*) are needed, our findings are consistent with photobiont turnover as an adaptation to shifting environmental conditions.

Although the sensitivity of lichen communities to pollution dominated research efforts in the 20th and early 21st century, we did not find a strong effect of heavy metals and other air-borne pollutants on our transplants (Supp. Fig 1). Although sites differed in pollutant exposure, this did not appear to impact the thallus-level photobiont communities over the duration of the experiment. This is consistent with research from other cities that have experienced significant improvements in air quality in recent decades, such that urban climate effects can affect lichen ecology as much or more than pollutants (Counoy *et al*. 2025; Koch *et al*. 2019; Munzi *et al*. 2014). This does not exclude additional differentiations between photobionts based on pollution tolerance, as was shown by Gasulla et al. (2025) in Madrid, but rather emphasizes the value of urban gradients to the study of future climate responses. Natural lichen communities, which reflect longer periods of exposure and local pools of available photobionts, might show differing patterns, and therefore require further study.

Symbiont turnover in response to environmental stress and change is also found in other photosynthetic symbioses such as corals. Coral photobiont composition has been shown to change directionally following warming (Buzzoni *et al*. 2026; Van Nynatten *et al*. 2025) and transplantation (Gantt *et al*. 2025), and introduction of warming adapted symbionts has even been suggested as a conservation strategy (Chan *et al*. 2025; Scharfenstein *et al*. 2024). While there are many biological differences between these symbioses–notably there is no evidence of lichen thallus “bleaching” through targeted expulsion of photosymbionts–, these similarities point to a more generalizable theory of symbiosis. This will require further studies of lichens and other symbioses to identify the conditions under which ecologically meaningful symbiont turnover is possible to better predict present and future vulnerabilities.

## Acknowledgments

Special thanks to all MSP-LTER (Minneapolis-St. Paul Metropolitan Area, Long Term Ecological Research Program) personnel, especially to Sarah Hobbie, Meredith Keller and Mary Marek-Spartz for providing valuable support for this work. Mary was also responsible for extracting and coding the buffer for NDVI and LST data. We thank all the land stewards who provided access to parks, as well as Caitlin Potter, Cedar Creek Ecosystem Science Reserve, from where our transplanted samples of *Flavoparmelia caperata* came from. We are grateful to all the field and laboratory assistants who helped with specimen collection, transplants and processing material in the lab (Ashley Darst, Adelaide Mahler, Mina Adabag and Samsam Hassan). We also thank Tami McDonald for helping with DNA extraction and amplification problem solving and Zan Tomko for helping organize outreach events. This work was possible thanks to the Minnesota Environment and Natural Resources Trust Fund as recommended by the Legislative-Citizen Commission on Minnesota Resources (LCCMR) grant 2023-152, who also funded the first author, and National Science Foundation funding (DEB-2045382) for the MSP-LTER. Part of this work was also funded by the Natural Resources Research Institute (NRRI), University of Minnesota (2018).

## Notes

### Competing Interest Statement

The authors have declared no competing interest.

