## Supplementary Table 1 and Figure 1 for "Moving to the city changed you! Rapid photobiont turnover enables acclimation of lichen symbioses to urban environments"

**Supplementary Material**

**Suppl. Table 1.** Results of the linear models between the physiological performance of the lichen thalli (gross and liquid assimilation, respiration and Fv/Fm) according to the OTU/Clade that predominates in their photobiont community (>70%) in relation to NDVI (Net Relatedness Index) in a buffer of 10m of the lichen transplants, and some metals: Fe, Pb and Zn). Gross and Net Carbon Assimilation, as well as Respiration were measured in μmol $m^{-2}$ $s^{-1}$.

| | | | | | | | | |
|---|---|---|---|---|---|---|---|---|
| **NDVI10** | | | | | | | | |
| | **A46** | | | | **I05** | | | |
| | Estimate | SE | t-test | P | Estimate | SE | t-test | P |
| Gross Carbon Assimilation | 7.30 | 3.09 | 2.37 | 0.03 | 1.21 | 3.29 | 0.37 | 0.72 |
| Net Carbon Assimilation | 8.51 | 3.29 | 2.59 | 0.02 | 0.66 | 3.68 | 0.18 | 0.86 |
| Respiration | 1.20 | 1.55 | 0.78 | 0.45 | -0.55 | 0.81 | -0.68 | 0.51 |
| Fv/Fm | 0.45 | 0.12 | 3.75 | 0.0007 | 0.10 | 0.14 | 0.67 | 0.51 |
| **Fe** | | | | | | | | |
| | **A46** | | | | **I05** | | | |
| | Estimate | SE | t-test | P | Estimate | SE | t-test | P |
| Gross Carbon Assimilation | -0.0007 | 0.002 | -0.31 | 0.76 | 0.0013 | 0.004 | 0.37 | 0.72 |
| Net Carbon Assimilation | -0.0018 | 0.002 | -0.75 | 0.46 | 0.0012 | 0.004 | 0.30 | 0.77 |
| Respiration | -0.0011 | 0.001 | -1.09 | 0.29 | -0.0001 | 0.0009 | -0.12 | 0.90 |
| Fv/Fm | -0.0001 | 0.0001 | -1.13 | 0.27 | 0.0002 | 0.0002 | 1.20 | 0.24 |
| **Pb** | | | | | | | | |
| | **A46** | | | | **I05** | | | |
| | Estimate | SE | t-test | P | Estimate | SE | t-test | P |
| Gross Carbon Assimilation | 0.009 | 0.24 | 0.04 | 0.97 | 0.24 | 0.44 | 0.55 | 0.59 |
| Net Carbon Assimilation | -0.09 | 0.26 | -0.33 | 0.74 | 0.25 | 0.49 | 0.52 | 0.61 |
| Respiration | -0.10 | 0.11 | -0.87 | 0.39 | 0.01 | 0.11 | 0.11 | 0.91 |
| Fv/Fm | -0.006 | 0.01 | -0.55 | 0.58 | 0.02 | 0.02 | 1.01 | 0.32 |
| **Zn** | | | | | | | | |
| | **A46** | | | | **I05** | | | |
| | Estimate | SE | t-test | P | Estimate | SE | t-test | P |
| Gross Carbon Assimilation | -0.01 | 0.02 | -0.82 | 0.42 | -0.01 | 0.07 | -0.17 | 0.87 |
| Net Carbon Assimilation | -0.02 | 0.02 | -1.14 | 0.26 | -0.04 | 0.08 | -0.46 | 0.65 |
| Respiration | -0.006 | 0.01 | -0.88 | 0.39 | -0.02 | 0.02 | -1.46 | 0.16 |
| Fv/Fm | -0.001 | 0.001 | -1.58 | 0.12 | -0.004 | 0.002 | -2.22 | 0.04 |

**Suppl. Figure 1.** Physiological performance metrics of *Flavoparmelia caperata* thalli according to the OTU/Clade that predominates in their photobiont community (>70% A46 or I05) with increasing Iron, Lead and Zinc thallus content. Points depict individual thalli that are colored based on this dominance. Metrics include **A-C.** gross carbon assimilation (μmol $m^{-2}$ $s^{-1}$), **D-F.** net carbon assimilation (μmol $m^{-2}$ $s^{-1}$), **G-I.** respiration (μmol $m^{-2}$ $s^{-1}$), and **J-L.** maximum photosynthetic efficiency of PSII (Fv/Fm).

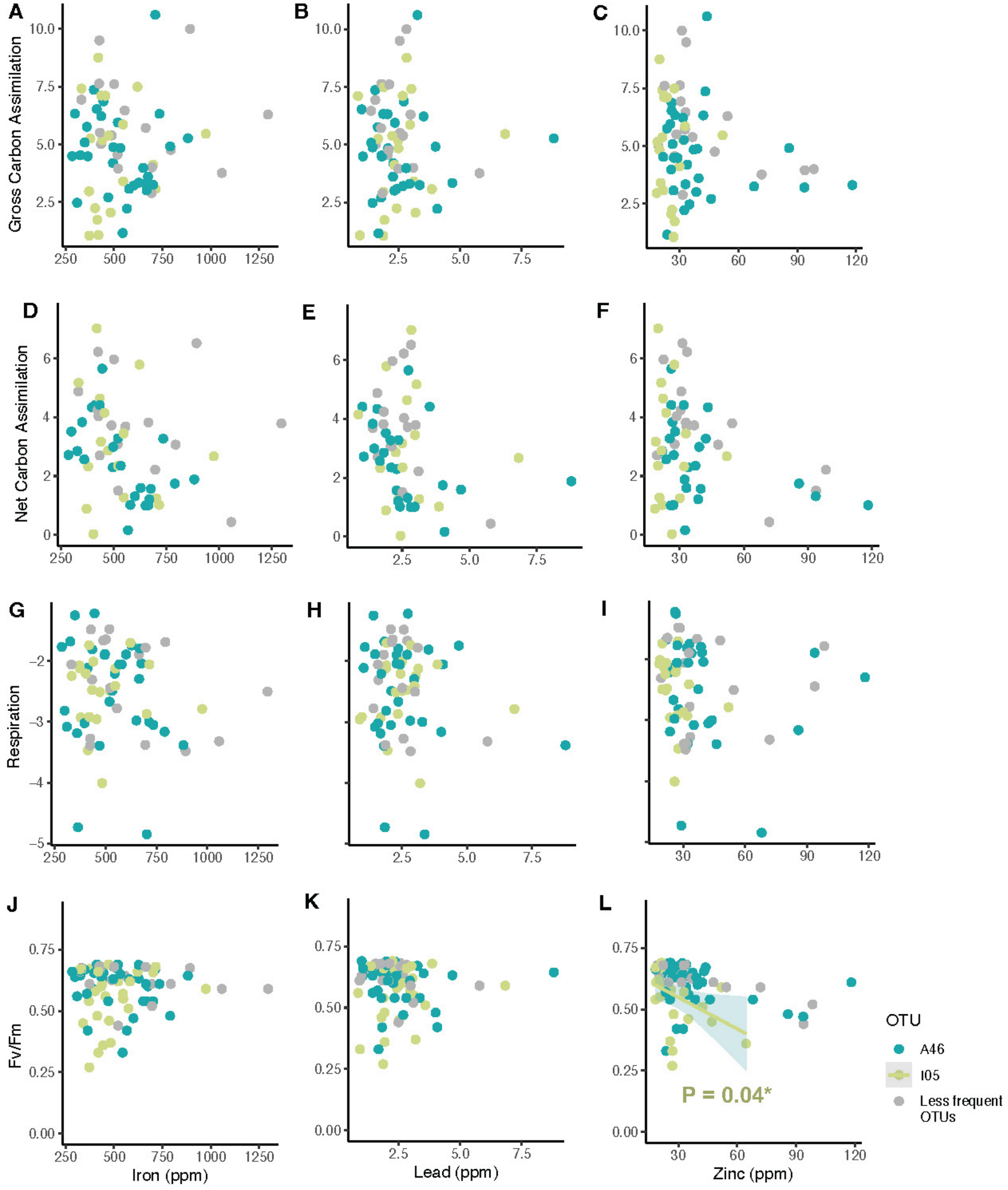
